# When the tap runs dry: The physiological effects of acute experimental dehydration in *Peromyscus eremicus*

**DOI:** 10.1101/2023.07.03.547568

**Authors:** Danielle M. Blumstein, Matthew D. MacManes

**Author notes:** (DMB, MDM.).

## Abstract

Desert organisms have evolved physiological, biochemical, and genomic mechanisms to survive the extreme aridity of desert environments. Studying desert-adapted species provides a unique opportunity to investigate the survival strategies employed by organisms in some of the harshest habitats on Earth. Two of the primary challenges faced in desert environments are maintaining water balance and thermoregulation. We collected data in a simulated desert environment and a captive colony of cactus mice (*Peromyscus eremicus*) and used lab-based experiments with real time physiological measurements to characterize the response to water-deprivation. Mice without access to water had significantly lower energy expenditures and in turn, reduced water loss compared to mice with access to water after the first 24 hours of the experiment. Additionally, we observed significant weight loss likely related to dehydration-associated anorexia a response to limit fluid loss by reducing waste and the solute load as well as allowing water reabsorption from the kidneys and gastrointestinal tract. Finally, we observed body temperature correlated with sex, with males without access to water maintaining body temperature when compared to hydrated males while body temperature decreased for females without access to water compared to hydrated, suggesting daily torpor in females.

## Introduction

Water is arguably the single most important factor for life on Earth and in organisms, water is stored in intracellular and extracellular spaces (Fitzsimons 1963). Dehydration occurs where there is a decrease in extracellular fluid volume caused when the loss is outpaced by fluid intake and metabolic water production, leading to a negative fluid balance and increased serum osmolality (Thomas et al. 2008). Regardless of the habitat, animals must regulate body fluids to protect against or cope with dehydration (Takei 2015). Mammals have developed many different mechanisms for body fluid regulation (Christian and Matson 1978; Frank 1988; Jirimutu et al. 2012; Marra et al. 2014; Yang et al. 2016; Kordonowy et al. 2017) and these mechanisms could aid in survival given the most well-supported climate change scenarios predict increased aridity (Mirzabaev et al., 2019).

Climate change is rapidly reshaping habitats globally and is predicted to continue (Hughes 2000; Parmesan and Yohe 2003; Parmesan 2006), modifying selective pressures for many populations (Hochachka and Somero 2002; Pörtner 2002; Pörtner and Farrell 2008). Understanding environmental tolerance and the capacity for adaptation in one species can provide insight into the potential for similar species to respond to increasingly extreme climatic patterns which are likely to affect many habitats. In recent years, many habitats have recorded some of the hottest temperatures to date (IPCC 2019; Stillman 2019), resulting in environmental thresholds that may exceed organismal tolerance. Furthermore, climate change is increasing global desertification rates, increasing water stress among wildlife (Loarie et al. 2009; Mirzabaev et al. 2019). To maintain viable populations, organisms must survive and successfully reproduce under climate warming and aridification by either using existing phenotypes and phenotypic plasticity, rapid evolution, or geographic range and phenological shifts (Hofmann and Todgham 2010; Hoffmann and Sgrò 2011; Brown et al. 2016). Despite climate change altering habitats and impacting populations, habitat distributions for rodents have remained remarkably stable over the last century of climate change, largely due to behavioral changes (Pardi et al. 2020; Riddell et al. 2021).

Mice of the genus *Peromyscus* have the widest distributions of any North American mammal and have unparalleled habitat diversity (Bedford and Hoekstra 2015). Several members of the genus, including the cactus mouse (*Peromyscus eremicus*) are native inhabitants of the arid deserts in southwest North America (Murie 1961; Pavlik 2008). Past studies have shown that cactus mice are extremely efficient at water retention (Kordonowy et al. 2017), have limited tissue damage when dehydrated (MacManes 2017), the slowest metabolism amongst the *Peromyscus* species (Mueller and Diamond 2001), have a suite of genomic adaptations (MacManes 2017; Tigano et al. 2020; Colella et al. 2021a), but lack the kidney modification present in kangaroo rats (Dewey et al. 1966; MacManes 2017). Furthermore, animals of this genus can be held in captivity (Crossland et al., 2014), have extensive genomic resources (Colella et al., 2021a; Tigano et al., 2020), and a wealth of samples collected historically and contemporaneously in natural history museums (Pergams and Lawler, 2009). Together, these features make the cactus mouse ideal for investigating water economy.

Desert habitats are characterized by such an extreme lack of precipitation which exerts a controlling effect on biological processes (Rocha et al., 2021). Daily temperatures in the Sonoran Desert can fluctuate by as much as 30-50 °C (Reid 1987; Sheppard et al. 2002). During the summer months, temperatures can reach upwards of 50 °C during the day, while at night they may drop to as low as 15 °C (Coppernoll-Houston and Potter 2018). In the winter, daytime temperatures are typically between 20 – 30°C, while nighttime temperatures can drop to near freezing (Boyd Deep Canyon Desert Research Center). Daily rainfall in the Sonoran Desert is relatively rare, with most areas receiving less than a centimeter of rain per year (Boyd Deep Canyon Desert Research Center). Organisms that are adapted to live in desert habitats must manage their water budgets over long dry and hot periods of time.

Here we expand on the long history of studies of organismal water management in desert taxa (Albright et al., 2017; Blumstein et al., 2022; Bradford, 1974; Cortés et al., 2000; Frank, 1988; Hayes et al., 1998; Kordonowy et al., 2017; MacMillen, 1983; Schmidt-Nielsen and Schmidt-Nielsen, 1952; Schmidt-Nielsen and Schmidt-Nielsen, 1952) to assess the physiological response to water deprivation in a hot and dry environment. To accomplish this, we compared multiple physiological responses, rate of water loss, energy expenditure, respiratory quotient, a suite of electrolytes, body weight, and body temperature, for mice with and without access to water for 72 hours to understand how animals survive the extreme head and aridity of deserts and further characterize *P. eremicus’* response to water deprivation.

## Methods

### Animal Care and Experimental Model

All animals used in this study were captive born, sexually mature, non-reproductive healthy adult male and female *P. eremicus*. Individuals were descended from wild caught animals from a dry-desert population in Arizona and maintained at the University of South Carolina Peromyscus Genetic Stock Center (Columbia, South Carolina, USA). Animal care procedures were approved by the University of New Hampshire Institutional Animal Care and Use Committee under protocol number 210602 and followed guidelines established by the American Society of Mammologists (Sikes and the Animal Care and Use Committee of the American Society of Mammalogists, 2016). Mice were housed in a large walk-in environmental chamber designed to simulate the temperature, humidity, and photoperiod of their native desert environment (Kordonowy et al. 2017; Colella et al. 2021b; Blumstein et al. 2022). The daytime (light) phase lasted for 12 hours (08:00 to 20:00) at a room temperature of 32°C and 10% RH followed by a one-hour transition period to the nighttime (dark) phase which lasted for 9 hours (21:00 to 06:00) at a room temperature of 24°C and 25% RH. To compete the cycle a two-hour transition period occurs to return the room to light phase conditions (Kordonowy et al. 2017; Colella et al. 2021b; Blumstein et al. 2022). Mice were provided a standard diet and fed *ad libitum* (LabDiet® 5015*, 26.101% fat, 19.752% protein, 54.148% carbohydrates, energy 15.02 kJ/g, food quotient [FQ] 0.89).

Prior to experimental conditions mice were weighed (rounded to the nearest tenth of a gram) on a digital scale. A temperature-sensing passive integrated transponder (PIT) tag (BioThermo13, accuracy ±0.02°C, BioMark®, Boise, ID, USA) was implanted subdermally between the shoulders of each rodent using a tag injector (Biomark® MK10). Animals were then allowed to recover individually in an experimental chamber for one week of observation before the experiments were started. Body temperature was recorded at noon and midnight via a Biomark® HPR Plus reader and weight was measured every noon over the duration of the experiment. A randomly selected set of animals were assigned to the two water treatment groups (n=9 of each treatment, female mice with water, female mice without water, male mice with water, and male mice without water, total n=36). At the start of the experiment (day 0, time 0, 10:00), water was removed from three of the chambers corresponding to those animals in the dehydration group. No mortality occurred during these experiments. Three days later, at the conclusion of the experiment, mice were euthanized via isoflurane overdose and decapitation, and we collected 120 µl of trunk blood for serum electrolyte measurement using an Abaxis i-STAT® Alinity machine. Using i-STAT CHEM8+ cartridges (Abbott Park, IL, USA, Abbott Point of Care Inc), we measured the concentration of sodium (Na, mmol/L), potassium (K, mmol/L), blood urea nitrogen (BUN, mmol/L), hematocrit (Hct, % PCV), ionized calcium (iCa, mmol/L), glucose (Glu, mmol/L), osmolality (mmol/L), hemoglobin (Hb, g/dl), chlorine (Cl, mEq/L), total CO_2_ (TCO_2_, mmol/L), and Anion gap (AnGap, mEq/L). Using Na, Glu, and BUN, we calculated serum osmolality. The experimental setup was repeated six times, three male batches and three female batches.

### Metabolic phenotyping

During the experiment mice were exposed to either water deprivation or normal conditions for three continuous days while being housed in transparent 9.5L respirometry chambers with dried cellulose-based bedding. Air was continuously pulled from the chambers using a pull flow-through respirometry system from Sable Systems International (SSI) starting with SS-4 Sub-Sampler Pumps, one for each chamber, at a rate of 1600 ml min_−1_ (96 l h_−1_). The SSI MUXSCAN was used to multiplexed air streams, measuring each chamber 120s approximately twice every hour. Finally, the Field Metabolic System (FMS, zeroed and spanned between each 72-hour experiment using dry gas with known concentrations of CO_2_ and O_2_) sub-sampled the airstream at 250 ml min^−1^ and measured water vapor, CO_2_, and O_2_ with no scrubbing.

### Calculations and Statistical Analysis

We analyzed our data using methods fully described in Colella et al. (2021b) and Blumstein et al. (2022). Rates of CO_2_ production, O_2_ consumption, and water loss were calculated using equations 10.6, 10.5, and 10.9, respectively, from Lighton (2018). Respiratory quotient (RQ, the ratio of VCO_2_ to VO_2_) and Energy expenditure (EE) kJ hr^-1^was calculated as in Lighton (2018, eq. 9.15). All downstream statistical analyses were conducted in R v 4.0.3 (R Core Team 2020). The R package mgcv::gamm was used and included the fixed effects; access to water and sex, and interacting nonlinear smoothing regression terms with pairwise fixed effect combinations as interactions; time in days and diurnal cycle (Lin and Zhang 1999; Wood 2017) and visualized using *gratia* (Simpson 2023). Experimental batches and the mice nested within the experimental batch were used as random effects to ensure we were not violating the assumption of independence. This allows us to explain the average differences between groups of mice instead of explaining differences between individual mice. To test for statistical significant (p < 0.05) differences in electrolytes after the treatments were applied and for each time point weight and body temperature were collected we used a student’s two-tailed t-test (stats::t.test) between the sexes for each experimental group.

## Results

### Rate of Water Loss

Both experimental groups, water access and water deprivation, had diurnal patterning of rate of water loss (RWL) with the highest occurring during the light phase and lowest during the dark phase (Figure 1A and 1 B). Each day of the experiment had similar patterns regardless of the treatment however, RWL was higher in males without access to water and for females lower or similar to the groups with water *ad lib* during day one of the experiment. For days two and three both males and females without access to water had lower RWL (Figure 1A and 1B).

**Figure 1.**
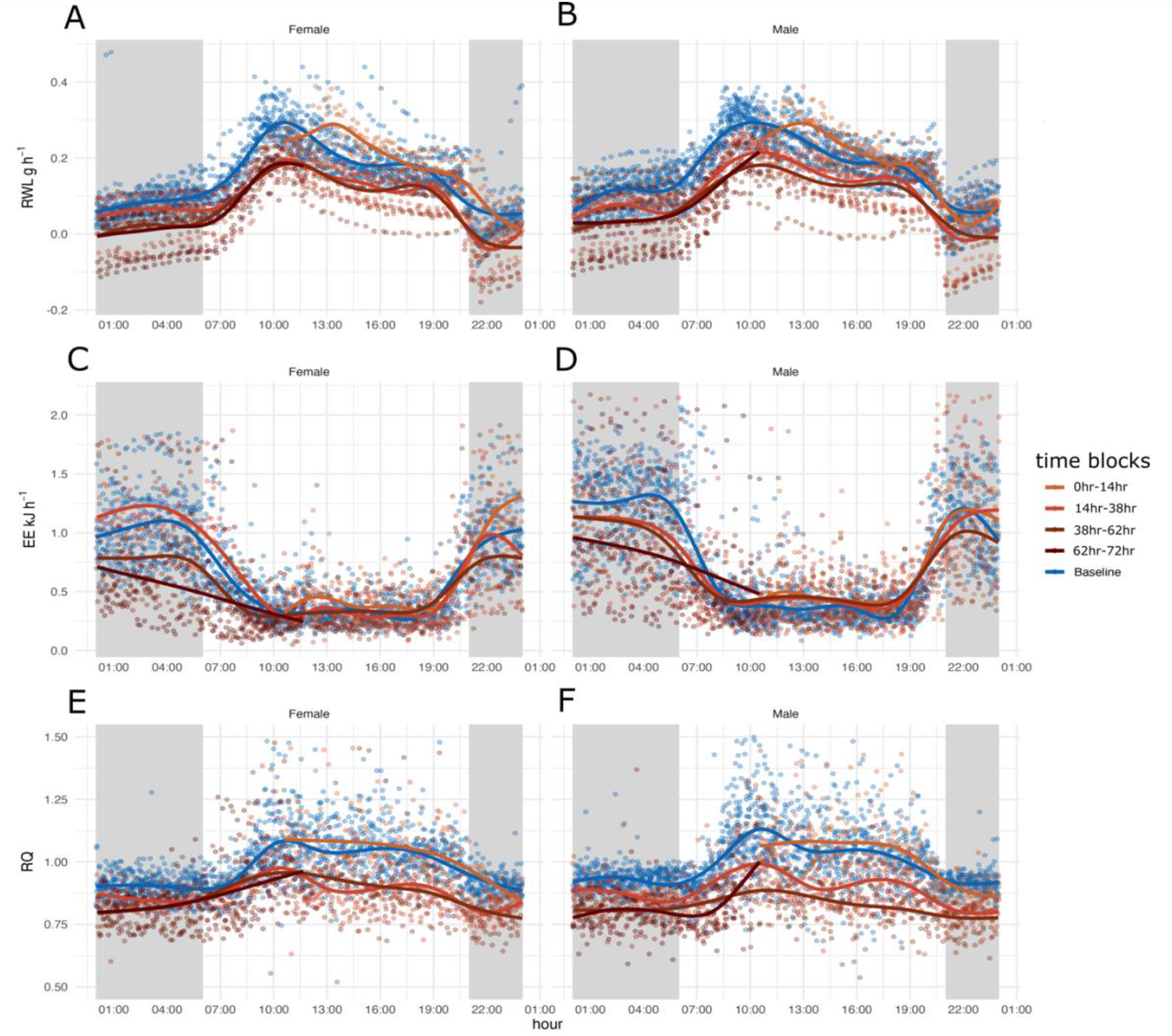
72 hours of respirometry data collection spit by sex for 18 adult males and 18 adult females plotted in a 24-hour window to display circadian patterns for each group: Baseline measurements of mice with water access (blue) and measurements of mice without access to water for one, two, three, and four days (four shades of brown). Shaded sections in gray indicate the dark phase when animals are active, and unshaded blocks indicate light phase when animals are inactive. A and B) 24-hour rate of water loss (RWL, H_2_O g hr^-1^) C and D) energy expenditure (EE kJ hr^-1^), and E and F) respiratory quotient (RQ), for females (A, C, E) and males (B, D, F).

Generalized additive modeling (GAM) analysis explained 77.2% of the deviance in RWL (Supplemental Figure 1, Supplemental Table 1). Significant predictors of RWL included sex (p < 2^-16^) and water access (p < 2^-16^) but not sex by water access (p = 0.13). All treatment combination splines were significant (Supplemental Table 1. For both males and females without access to water, the curves for time in days by 24-hour cycle were very complex, oscillating with the light dark cycle and decreasing over time (Figure 1A and 1B). The curves for time in days for males and females with access to water oscillated with the light dark cycle as well (Figure 1A and 1B). Generally, mice without water had higher RWL during the first 24 hours and lower RWL for the remainder of the experiment based on GAM analysis and visualization (Supplemental Figure 1, Supplemental Table 1, Figure 1A and 1B). When comparing the four curves (males without water, males with water, females without water, females with water), RWL was similar to mice with access to water converged during the light to dark transition phases, with the exception of the first transition (Supplemental Figure 1).

### Energy Expenditure

Males and females in both experimental groups, water access and water deprivation, show diurnal pattering, with the highest EE occurring during the dark (active) phase and the lowest EE occurring during the light (inactive) phase (Figure 1C and 1D). Each day of the experiment for males and females with and without water has a similar pattern, elevated during the dark phase, and reduced during the light phase, regardless of the treatment.

During the first 24 hours, EE was highest for females without access compared to males without access to water and all mice with access to water. EE decreased over the subsequent 48 hours for mice without access to water with females without access to water having the lowest EE compared to males without access to water and all mice with access to water during the dark phase of days two and three of the experiment (Supplemental Figure 2, Supplemental Table 2, Figure 1C and 1D). Mice were manually weighted at 12:00 every day, resulting in a transient increase of EE at that time (Figure 1C and 1D). The GAM analysis explained 61.4% of the deviance in EE with significant predictors being sex (p < 2_-16_), water access (p < 2_-16_), and sex * water access (p = 0.0464). All treatment combination splines were significant (Supplemental Table 2).

### Respiratory Quotient

RQ had diurnal patterning for both experimental groups and for both sexes (Figure 1E and 1F). RQ was highest during the light phases (Figure 1E and 1F) and lowest and comparable to the FQ during the dark phases (Figure 1E and 1F) 43.3% of the deviance was explained in the GAM analysis with significant predictors being sex (p < 2_-16_), water access (p = 4.14_-08_), and the interaction between sex and access to water (p = 1.41_-07_). All treatment combination splines were significant (Supplemental table 3) and complex, oscillating with the light dark cycle (Figure 1E and 1F, Supplemental Figure 3). Males and females without water access had higher RQ compared to mice with water access during the first 24-hours based on GAM analysis and visualization (Supplemental Figure 3, Supplemental Table 3). Interestingly, males without access to water had the lowest RQ of any group over the course of the entire experiment during the second dark phase and for the remainder of the experiment (Supplemental Figure 3).

### Electrolytes, Weight, and Body Temperature

Several electrolytes were significantly different when comparing males with and without access to water and females with and without access to water (male and female Na p = 0.0016 and p = 0.0026 respectively, BUN p = 0.001/0.003, Hct p = 0.002/0.001, osmolality p = 8.2^-05^/0.0001, Cl p = 0.02/0.007, Hb p = 0.017/0.009, and TCO_2_ female p = 0.017) (Figure 2). No electrolytes were significantly different when comparing males to females for either water treatment (Figure 2).

**Figure 2.**
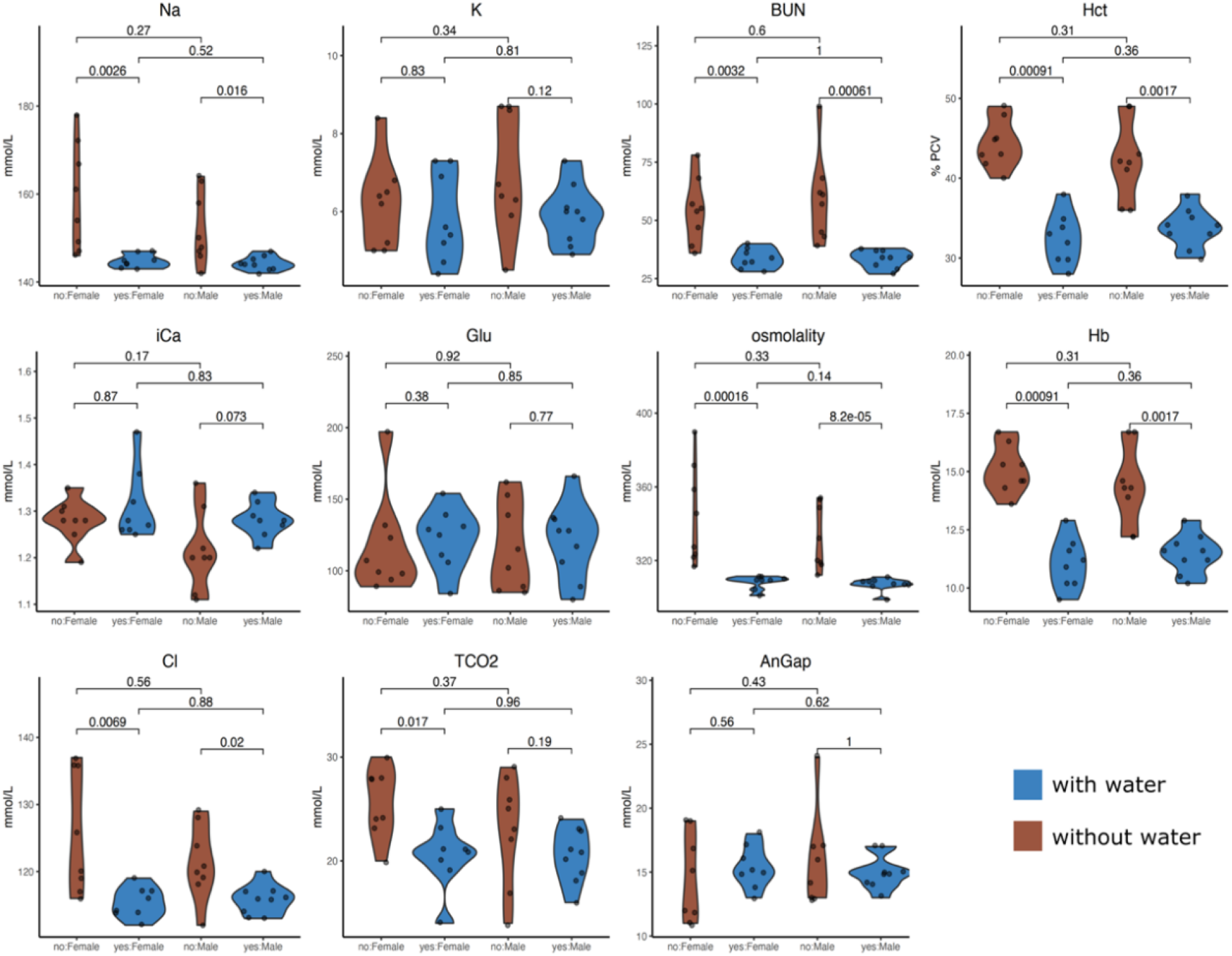
Violin plots showing the distribution of serum electrolyte measurements (Na = sodium (mmol/L), K = potassium (mmol/L), BUN = blood urea nitrogen (mmol/L), Hct = hematocrit (% PCV), iCa = ionized calcium (mmol/L), Glu = glucose (mmol/L), osmolality (mmol/L), Hb = hemoglobin (g/dl), Cl = chlorine (mEq/L), TCO_2_ = total CO_2_ (mmol/L), and AnGap = Anion gap (mEq/L), for female and male *Peromyscus eremicus* with (blue) or without (brown) access to water for 72 hours. Observations (n=9 of each treatment, total n=36) are represented by black dots. P-values from pairwise t-tests are reported above the brackets.

While the weights of males and females where insignificant at the beginning of the experiment, both sexes lost weight over the course of the water deprivation experiment with the most weight loss occurring in the first 24 hours without water (Figure 3). When comparing males without access to water to males with water access and females without access to water to females with water access, mice without water weighted significantly less then mice with water at 24 hours (p = 0.024, 0.019), 48 hours (p = 0.004, 0.002) and 72 hours (p = 0. 001, 0.005) (Figure 3A and 3B). Only animals held without water lost weight (Figure 3C and 3D), and analysis of these changes were significantly different at all timepoints after water had been removed (24 hours, p = 4.1_-05_, 4.1_-05_), (48 hours, p = 0.001, 4.1_-05_), and (72 hours, p = 4.1_-05_, 0.001).

**Figure 3.**
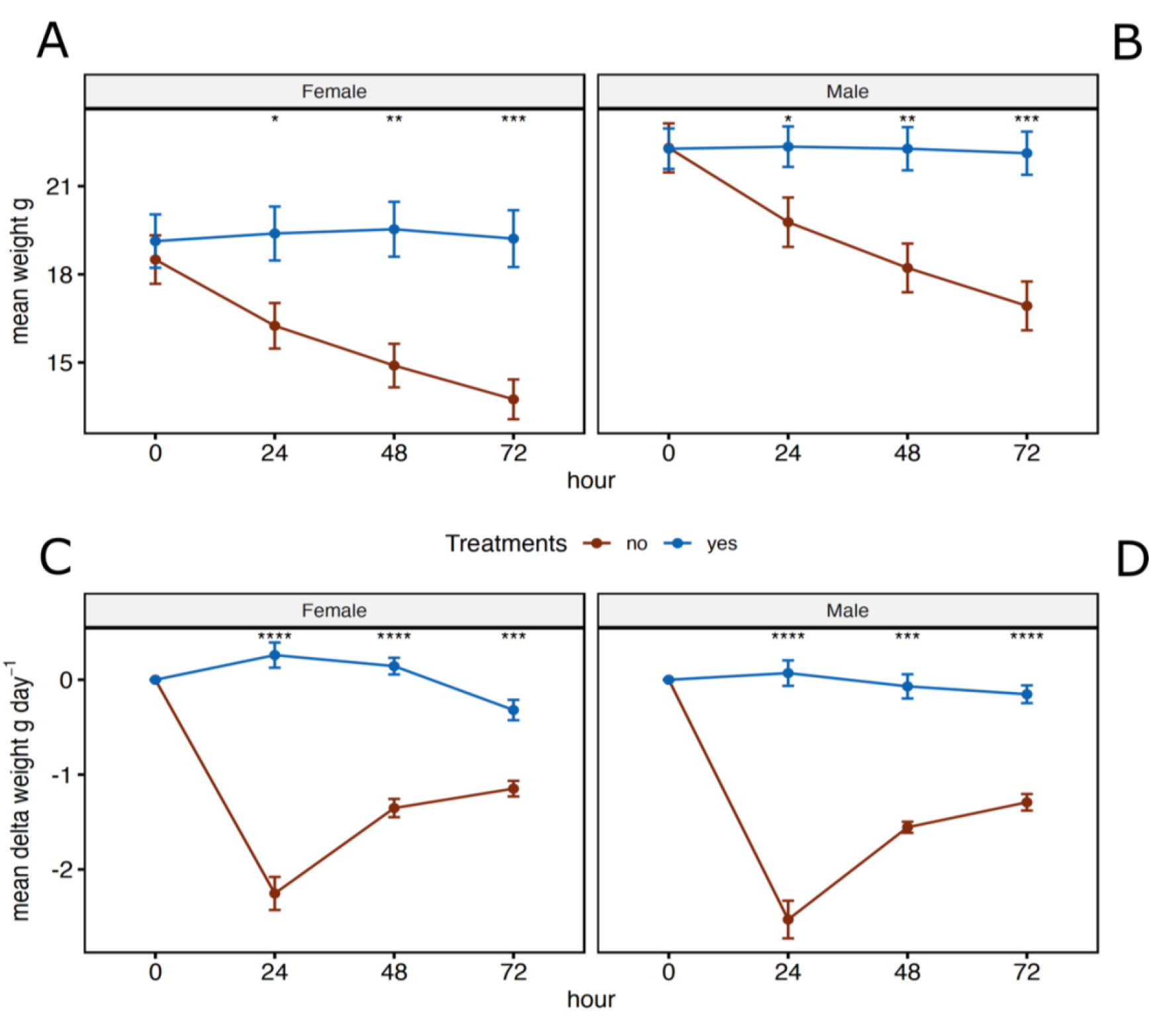
Mean weights (A and B) and mean delta weights (C and D) for female and male *Peromyscus eremicus* with (blue) or without (brown) access to water every 24 hours over the course of the 72-hour experiment. Error bars represent the standard error and * across the top denote statical significance from t-tests between the two treatments, with and without water, at each timepoint (* p <= 0.05, **: p <= 0.01, ***: p <= 0.001, ****: p <= 0.0001).

Body temperature showed diurnal pattering with the highest body temperature during the dark (active) phase and the lowest during the light phase (**Figure 4**. For females, body temperature followed a similar pattern as described above and were significantly lower for mice without access to water at 24 hours (p = 0. 001), 36 hours (p = 0.005), 48 hours (p = 0.001), 60 hours (p = 0.002), and 72 hours (p = 0.0003) while males were not significantly different at any of the time points (**Figure 4**).

**Figure 4.**
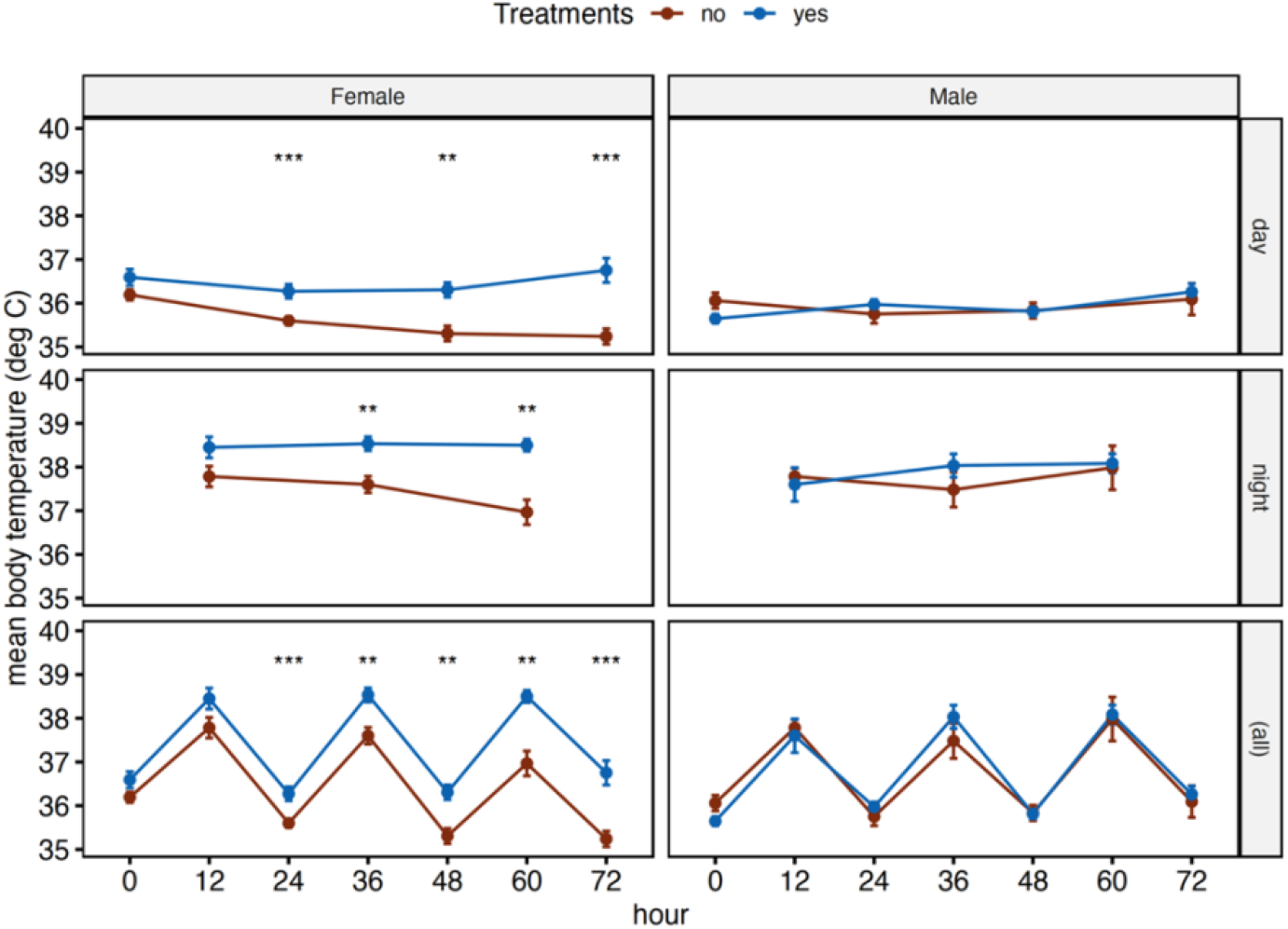
Mean body temperatures for female and male *Peromyscus eremicus* with (blue) or without (brown) access to water every 12 hours over the course of the 72-hour experiment. The top row of graphs are measurements taken only during the light phase, middle row are measurements taken only during the dark phase, and bottom row represents all the measurements. Error bars represent the standard error and * across the top denote statical significance from t-tests between the two treatments, with and without water, at each timepoint (* p <= 0.05, **: p <= 0.01, ***: p <= 0.001, ****: p <= 0.0001).

## Discussion

Physiological mechanisms can act as a buffer, expanding organismal tolerance to new or extreme environments (Bijlsma and Loeschcke, 2005; Gabriel, 2005; Lau et al., 2017; Lui et al., 2015; Wilson and Franklin, 2002), however, biochemical and physical constraints eventually limit physiological capacity (Campbell-Staton et al., 2021; Velotta and Cheviron, 2018; Velotta et al., 2018), and ultimately determine a population’s persistence (Parmesan, 2006; Parmesan and Yohe, 2003; Van der Putten et al., 2010). In xeric habitats, further increased ambient temperature is compounded by reduced water availability, potentially affecting an animal’s ability to maintain homeostasis of body temperature and fluids (Reece et al., 2015). Given the current rapid pace of climate change (IPCC 2019), it is vital that we understand how species are responding to changes in their environment. Increased drought and changes in precipitation patterns are having several impacts on the availability of water, both in terms of availability, quantity, and quality (IPCC, 2019; Mirzabaev et al., 2019).

Organisms maintain water homeostasis in several ways, including seeking out sources of free-flowing water, preformed dietary water (Frank 1988; Orr et al. 2015) and/or water produced by metabolism. However, if adequate water is not acquired, dehydration can negatively affect an animal’s ability to regulate its body temperature, impair its’ cardiovascular function, and decrease perfusion to organ systems. Specifically, dehydration results in a decrease in blood volume and increase in osmolality, primarily driven by the increase in serum sodium levels (Leib et al., 2016; Thornton, 2010). As a result, several neurohormonal systems are activated to maintain blood pressure to perfuse tissues appropriately (Kaufmann et al., 2020). Water is recovered in the gastrointestinal tract (Thiagarajah and Verkman, 2018) and in the kidneys it is reabsorbed from the tubule system back into the blood stream (Fuller et al., 2020; Kortenoeven and Fenton, 2014). In severe cases, dehydration can lead to organ failure and death.

We explored the relationships and tradeoffs between thermoregulation, osmoregulation, and energy expenditure, of desert adapted mice without access to drinking water for three days while housed in an environmental chamber that simulated the desert environment. There are multiple avenues of water loss, including loss via urine and feces, as well as via cutaneous evaporation and respiration, and the measurements presented here represent their sum. We observed that when water was removed, energy expenditure and evaporative water loss are reduced in both sexes (presumably to conserve body water) at the expense of homeothermy, resulting in lower core body temperature in females but not in males. Though it may save water and/or energy, these physiological shifts could ultimately increase the risk of mortality and decrease fitness if water continues to be unavailable for extended periods of time.

### Weight loss and water deprivation

Our study targeted responses to water deprivation, investigating how physiological variables changed in response to dehydration throughout the course of the experiment. We found that in response to water deprivation, cactus mouse phenotypic responses changed rapidly. During the first 24-hours of the water-deprivation experiment both males and females increased EE, resulting in an increase in RWL, and a significant decrease in body weight. The reasons behind this dramatic shift are unclear but may be a result of 1) a behavioral response related to searching for drinking water and or 2) suppression of food intake as suggested by pilot studies.

The relationship between eating and drinking has been extensively studied (Fitzsimons and Le Magnen, 1969; Kissileff, 1969; Smith, 2000; Watts, 1998; Zorrilla et al., 2005) and it has been documented that dehydration-anorexia that is an adaptive response to limit fluid loss (Watts and Boyle, 2010), as typically the processing of food requires the use of water. Previous studies have demonstrated that dehydrated animals with *ad lib* food match the same attributes of food restricted animals, such as expression of hypothalamic neuropeptide genes, leptin and insulin levels, and corticosterone concentrations (Watts et al., 1999). Furthermore, the reduction of food intake results in a series of adaptive responses that target GI function, allowing for the absorption of the osmotically sequestered water that is normally in the GI (Kutscher, 1966; Lepkovsky et al., 1957; Schoorlemmer and Evered, 1993). Finally, reduced food intake also reduces the solute load (Rowland, 2007) and the need for removal of waste products via urinary water loss (Schoorlemmer and Evered, 1993). In the study discussed herein, several tissues, including the GI tract, were removed at the conclusion of the experiment for future RNAseq analysis, and the GI tract was empty of food and feces (unpublished observations), suggesting that the intake of solid food had been decreased dramatically. As mentioned above, we saw a significant decrease in weight during the first 24 hours of the experiment, however, the RWL during the first 24 hours is not enough to account for the weight loss, suggesting weight loss through other means such as dehydration-anorexia (Armstrong et al., 1980; Hamilton and Flaherty, 1973).

Previous studies have found that access to water (Hochman and Kotler 2006; Shrader et al. 2008; Levy et al. 2016) and specific dietary composition (Blumstein et al., 2022; Frank, 1988; Manlick et al., 2021; Orr et al., 2015; Schmidt-Nielsen, 1975; Schmidt-Nielsen and Adolph, 1964; Wolf and del Rio, 2003) strongly effects populations living in arid environments. These external factors influence species distributions (McKee et al. 2015), modifying foraging decisions (Gedir et al. 2016, 2020), and altering behavior and reproduction (Douglas 2001; McKinney et al. 2001; Cain et al. 2008). In the wild, cactus mice have been documented shifting diet seasonally, consuming arthropods during the winter (Hope and Parmenter, 2007), and transitioning to the consumption of cactus seeds and/or fruits during the summer (Hope and Parmenter, 2007; Orr et al., 2015). In addition to preformed water, the composition of diet is also very important for *P. eremicus* as described in Blumstein et al. (2022). Specifically, mice fed a diet low in fat with *ad lib* water lost significantly more water and had electrolyte levels suggesting dehydration compared to mice fed a diet higher in fat, suggesting a limited capacity to tolerate water deprivation if optimal foods become less abundant (Blumstein et al. 2022). Furthermore, the temperatures required to balance evaporative water loss with metabolic water production on dry seed are much lower than what occurs during the summers in desert regions (MacMillen and Hinds 1983; Walsberg 2000), suggesting that *P. eremicus* may not be able to survive on a only a dry diet, unlike the Heteromyids, which survive on dry diets alone (Frank, 1988; Schmidt-nielsen et al., 1948).

Consistent with predictions of altered physiology and behavior mediated by water restriction, we recorded a decrease in EE, RWL, body weight, and body temperature and a shift in serum electrolytes in water deprived *P. eremicus* during all three 24-hour time blocks. While males and females without access to water had different magnitudes of change in EE and RWL throughout the duration of the study, both metabolic rates and the rate at which water is lost decreased, similar to what has been recorded in other desert organisms (Schmidt-Nielsen et al. 1967; Taylor 1969; Finch and King 1982). EE and RWL are inherently related in animals as lower EE leads to lower water loss by decreasing the amount of dry air passing through the respiratory track (McFarlane and Howard 1972). Furthermore, catabolism of different diets vary in the amount of available energy (Sánchez-Peña et al., 2017), water potential, as well as their obligatory water loss (Schmidt-Nielsen, 1975). At lower humidity, oxidation of carbohydrates produces a net metabolic water gain while lipids and proteins result in water loss, mainly through urination which is required to remove products of their metabolism like urea.

### Sexual Dimorphism

Interestingly, males and females responded differently to lack of water, with body temperature being the most notable difference. Females decreased their body temperature while males maintained their body temperature when compared to their hydrated counterparts. Whether this sexually dimorphic response is a strategy or consequence is an open-ended question that cannot be answered using the data presented here, this response may be the product of high costs of reproduction in females, but not males. Indeed, similar patterns of sexual dimorphism in response to resource availability has been observed in other rodent species (Cranford 1977; Randolph et al. 1977; Murray and Smith 2012). Previous studies hypothesized that sexual dimorphism differences can be explained by differences in body size, metabolism, respiratory rate, or activity (Cryan and Wolf, 2003). While we do not have direct measurements of respiratory rate or activity, the production of CO_2_ follows the patterns of EE, providing indirect yet strong evidence that respiratory and metabolic rates (EE) as well as activity are all sexually dimorphic, consistent with observations in humans (Glucksmann 1974; Mittendorfer 2005; Pomatto et al. 2018) and has also been observed in *P. eremicus* by McNab and Morrison (1963) and Colella et al. (2021b).

Male reproduction is mainly limited by access to females (Bateman 1948), therefore, torpor or estivation by males could reduce male reproductive success. Furthermore, sperm quantity and quality is dependent on body temperature (Moore, 1926; Pérez-Crespo et al., 2008) and while typically resolved by externalizing the testes to the scrotum during excessive heat, a decrease in body temperature, as is seen in females (discussed below), could reduce sperm viability. Maintaining consistent body temperatures also allows for regular biological reactions, such as enzymatic processes and protein folding which have evolved to function best at a single temperature and can influence a series of functions not directly related to reproduction, such as growth rate, metabolic biorhythms, and environmental sensing (Glucksmann 1974; Hochachka and Somero 2002; McPherson and Chenoweth 2012; Calisi et al. 2018). Our data supports this as body temperature was unchanged for dehydrated males compared to their hydrated counterparts for the entire experiment, suggesting a maintenance of reproductive investment at the cost of long-term survival.

In contrast, female reproduction is primarily limited by their access to resources (Bateman 1948), in this case water. During the course of our study, female body temperature, EE, and RWL all decreased, suggesting torpor and or estivation, consistent with MacMillen (1983). Specifically, homeostatic responses such as adaptive heterothermy, a process which reduces evaporation by storing body heat, reduces the air to body temperature gradient thus decreasing inward heat flow, minimizes water loss from evaporative cooling (Schmidt-Nielsen et al. 1956; Schoen 1972; Taylor 1972; Cain et al. 2008), and in small endotherms with high surface area to volume ratios heterothermy can lead to substantial energy and water savings (Walsberg 2000; Speakman and Król 2010; Turbill and Stojanovski 2018). For females, reproductive demands are especially high, particularly during pregnancy and in lactating females (not measured in this study, Sorensen et al. 2005; Murray and Smith 2012), and minimizing energy costs or allocating pulses of resources to reproductive energy could increase reproductive success (Smith et al. 2014; Flores-Manzanero et al. 2019). While homeostatic responses are quite common among endotherms (Boyles et al. 2011, 2013; Canale et al. 2012; McGuire et al. 2014; Dammhahn et al. 2017) and are essential for short term survival, they incur energetic, resource, and or fitness costs when the disturbance lasts longer than the homeostatic tolerance (Wingfield et al. 1992; Boonstra 2004; Canale et al. 2012; McGuire et al. 2014; Dammhahn et al. 2017).

### Electrolytes

In order to gain a deeper understanding of how water deprivation affects the physiological functioning of endotherms in desert environments, we collected serum electrolyte data from both males and females with and without access to water at the end of the experimental period. Electrolytes are essential for all physiological functions, including regulating fluid balance, transmitting nerve impulses, and maintaining the acid-base balance (Hasona and Elasbali, 2016). Additionally, electrolyte levels can provide insight into an individual’s overall metabolic state, renal function, and can be indicative of dehydration, kidney disease, or heart failure (Kutscher 1968; Cheuvront et al. 2010).

The kidney typically ensures that fluid and electrolyte balance remain within a narrow range, and this is conducive to efficient biochemical and physiological processes. Altered electrolytes, such as K, iCa, and Na, are associated with dehydration (Abubakar and Sule, 2010; Cheuvront et al., 2010; Kutscher, 1966), and may result in fatigue, cognitive disfunction, and changes in osmotic pressure which may affect blood pressure. More severe electrolyte abnormalities may cause cardiac arrhythmias, and lead to death (Abubakar and Sule, 2010). We uncovered significant differences in electrolyte values between water treatments (Na, BUN, Hct, osmolality, Hb, Cl, and total CO_2_), suggestive of dehydration, but synthetic markers of renal function were unchanged. Together, supporting findings from (Kordonowy et al., 2017), this suggests that end-organ perfusion is maintained despite dehydration.

Despite being statistically insignificant, glucose trended downwards for males and females without water when compared to their hydrated counterparts. During fasting, blood glucose levels decrease due to a lack of glucose absorbed from the GI tract (Jensen et al., 2013). Previous studies have shown that the duration of fasting significantly affects blood glucose levels up to 72 hours, but after 72 hours there is no further decrease (Jensen et al., 2013). In humans, glucose concentrations are maintained regardless of the duration of starvation (Watford, 2015). Initially, carbohydrates are depleted during the first 24 hours, however, during prolonged starvation gluconeogenesis provides glucose by breaking down skeletal muscle proteins (Watford, 2015). While the current study does not measure food intake, glucose is being maintained, suggesting they could be reducing in food intake consistent with other studies of dehydration-anorexia in rodents and or shift toward increased glycogenolysis and lipolysis to maintain glucose concentrations (Salter and Watts, 2003; Schoorlemmer and Evered, 2002; Watts and Boyle, 2010), meaning the liver is possibly serving as a buffer for blood glucose concentration.

## Conclusion

The extreme aridity of desert environments plays a role in shaping biological processes however, the physiological mechanisms that allow animals to maintain salt and water homeostasis are still not well understood. Rapid climate change can challenge this tolerance. The cactus mouse (*Peromyscus eremicus*) is native to the arid deserts in southwest North America. Past studies have shown that cactus mice are highly adapted to desert conditions, with efficient water retention and dehydration tolerance. Therefore, cactus mice represent an interesting experimental model to examine physiological adaptations and thresholds.

In this study, we explore the physiological mechanisms that enable cactus mice to survive in desert habitats. By integrating laboratory-based experiments with long-term physiological data collected from a captive colony of cactus mice in a simulated desert environment, we investigate their response to water deprivation. Our findings reveal that mice without access to water exhibit significantly lower energy expenditures, leading to reduced water loss compared to mice with access to water. Moreover, significant weight loss was observed during the first 24 hours, likely attributed to dehydration anorexia— an adaptive response aimed at limiting fluid loss by reducing waste and the solute load, while facilitating water reabsorption from the kidneys and gastrointestinal tract. Furthermore, our observations indicate a relationship between body temperature and sex. Males without access to water maintained their body temperature compared to hydrated males, while females without access to water experienced decreased body temperature, suggesting the occurrence of daily torpor in females as an adaptive response, likely related to reproductive investment.

By examining the physiological responses of water deprived *P. eremicus*, we gain valuable insights into how adaptations developed over long evolutionary timescales. Given the current global climate change and the escalating desertification trends, it becomes imperative to investigate the plasticity and mechanisms of response in desert-adapted species. Such investigations hold the potential to enhance our understanding of organismal responses to the increasingly unpredictable climatic conditions.

## Acknowledgments

We would like to thank members of the MacManes lab including Ella Caughran, Molly Kephart, Sean Pierre-Louis, Sarah Nicholls for helpful comments and support on previous versions of the manuscript. The Animal Resources Office and veterinary care staff at the University of New Hampshire for colony maintenance and care. This work was supported by the National Institute of Health National Institute of General Medical Sciences (R35 GM128843 to M.D.M.).

## Author Contributions

Conceptualization: M.D.M.; Methodology: D.M.B., M.D.M.; Formal analysis: D.M.B., Investigation: D.M.B., Resources: M.D.M.; Writing - original draft: D.M.B.; Writing - review & editing: D.M.B., M.D.M.; Visualization: D.M.B; Supervision: M.D.M.; Project administration: M.D.M.; Funding acquisition: M.D.M.

## Data Availability

Macro processing files, processed respirometry data, and cage sampling scheme files are available on Zenodo: https://zenodo.org/record/8091766. All R scripts used in this project are available through GitHub at: https://github.com/DaniBlumstein/dehy_phys.

## Supplemental

**Figure 1.**
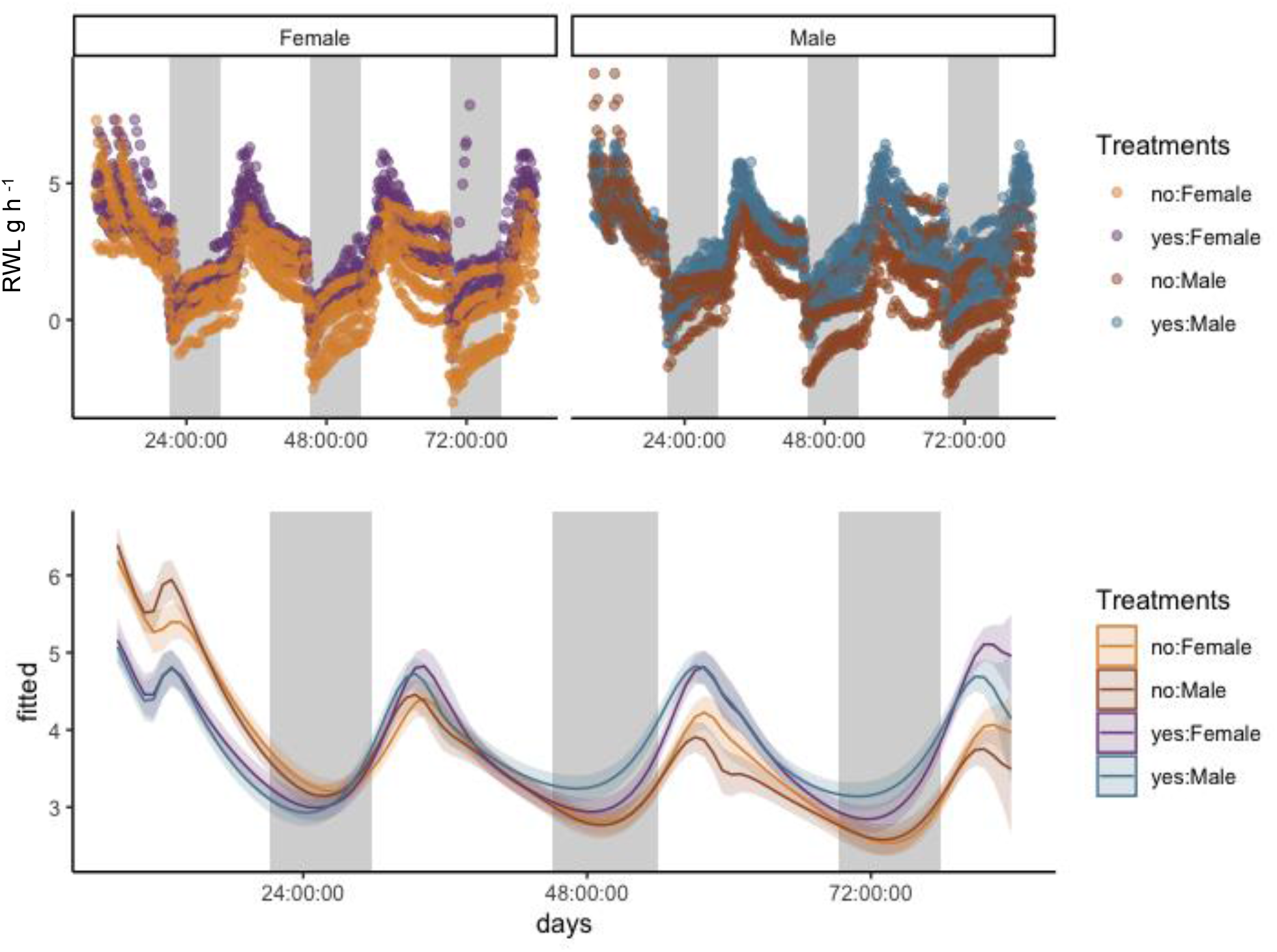
Raw plotted data (top) and general additive mixed models (GAMM) graph (lower) for rate of water loss (RWL g h_-1_) for female and male *Peromyscus eremicus* with and without access to water. The smoothing curves for each response variable included two fixed effects; water treatment (yes vs no) and sex, two random effects; mouse identification number and date of data collection, and two regression terms: time in days and diurnal cycle. For the lower graph, the y-axis is the effect of the x-axis on RWL as estimated by a multivariable GAMM. Shaded areas are 95% confidence intervals.

**Figure 2.**
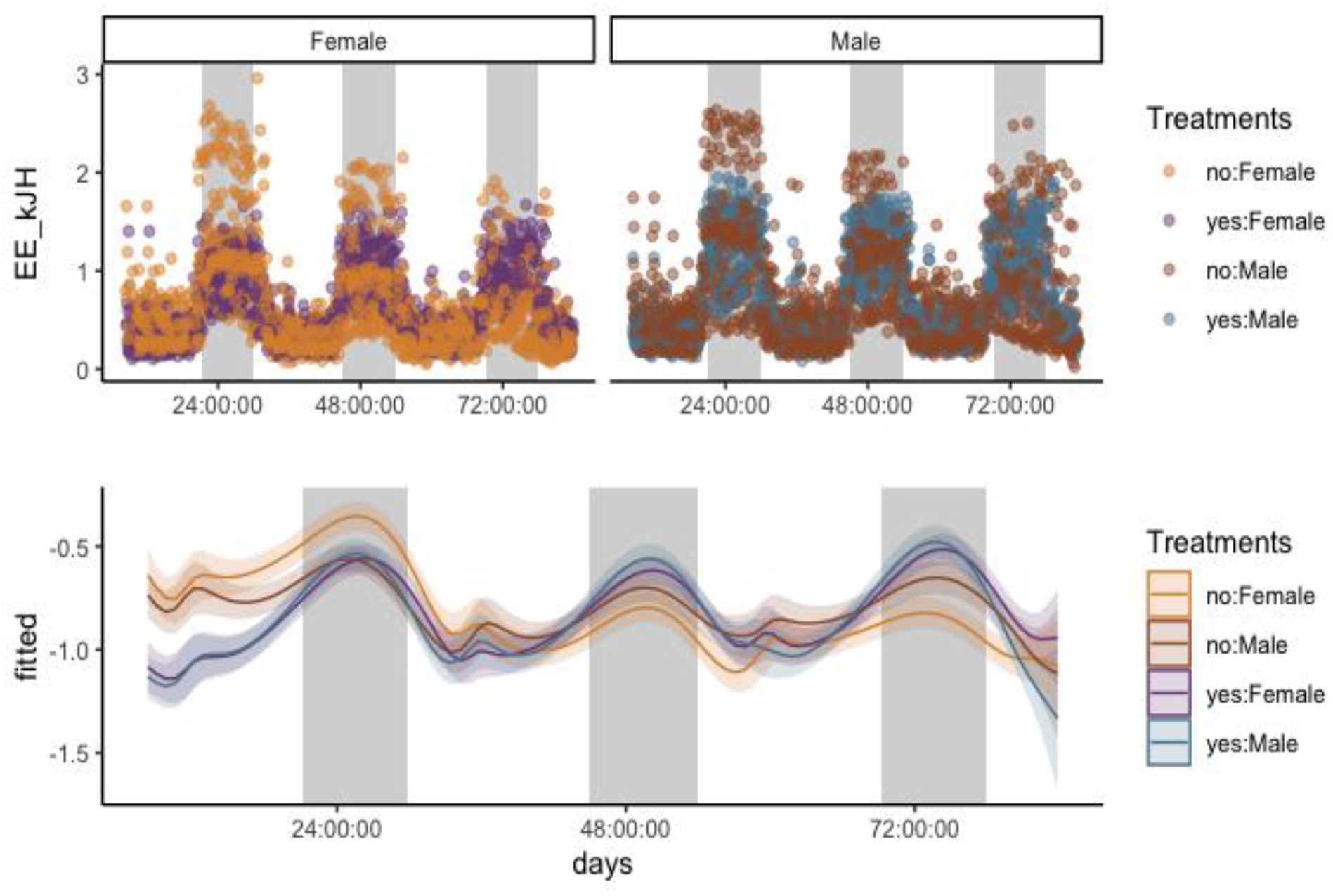
Raw plotted data (top) and general additive mixed models (GAMM) graph (lower) for energy expenditure (EE kJ h_-1_) for female and male *Peromyscus eremicus* with and without access to water. The smoothing curves for each response variable included two fixed effects; water treatment (yes vs no) and sex, two random effects; mouse identification number and date of data collection, and two regression terms: time in days and diurnal cycle. For the lower graph, the y-axis is the effect of the x-axis on EE as estimated by a multivariable GAMM. Shaded areas are 95% confidence intervals.

**Figure 3.**
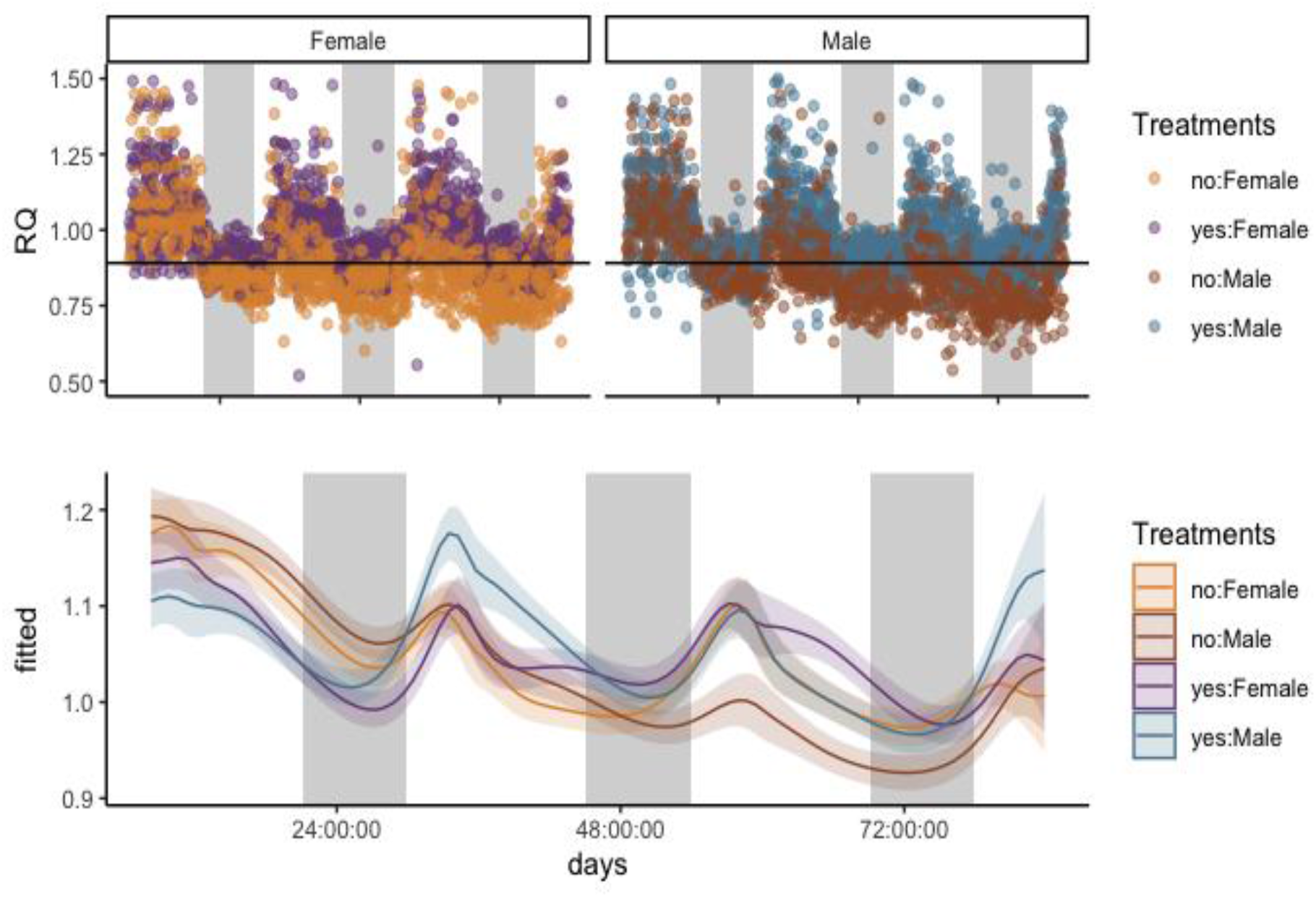
Raw plotted data (top) and general additive mixed models (GAMM) graph (lower) for Respiratory Quotient (RQ) for female and male *Peromyscus eremicus* with and without access to water. The smoothing curves for each response variable included two fixed effects; water treatment (yes vs no) and sex, two random effects; mouse identification number and date of data collection, and two regression terms: time in days and diurnal cycle. For the lower graph, the y-axis is the effect of the x-axis on RQ as estimated by a multivariable GAMM. Shaded areas are 95% confidence intervals.

**Table 1.** Generalized additive mixed models (GAMM) statical model and results for rate of water loss (RWL, H_2_O g/hr^-1^).

|  | Estimate | Std. Error | t value | Pr(>\|t\|) |
| --- | --- | --- | --- | --- |
| (Intercept) | 1.67969 | 0.02029 | 82.775 | <2e-16 *** |
| H2Oyes | 0.83920 | 0.02874 | 29.195 | <2e-16 *** |
| SexMale | 0.27509 | 0.02878 | 9.557 | <2e-16 *** |
| H2Oyes:SexMale | 0.06247 | 0.04054 | 1.541 | 0.123 |

|  | edf | Ref.df | F | p-value |
| --- | --- | --- | --- | --- |
| s(days,time_in_D):ttno:Female | 56.81 | 71.02 | 67.57 | <2e-16 *** |
| s(days,time_in_D):ttyes:Female | 63.72 | 78.17 | 55.76 | <2e-16 *** |
| s(days,time_in_D):ttno:Male | 62.10 | 76.51 | 66.08 | <2e-16 *** |
| s(days,time_in_D):ttyes:Male | 62.61 | 77.02 | 58.38 | <2e-16 *** |

Signif. codes: 0 '\*\*\*' 0.001 '\*\*' 0.01 '\*' 0.05 '.' 0.1 ' ' 1
R-sq.(adj) = 0.764 Deviance explained = 77.3%
-REML = 8209.6 Scale est. = 0.65701 n = 6463

**Table 2.**
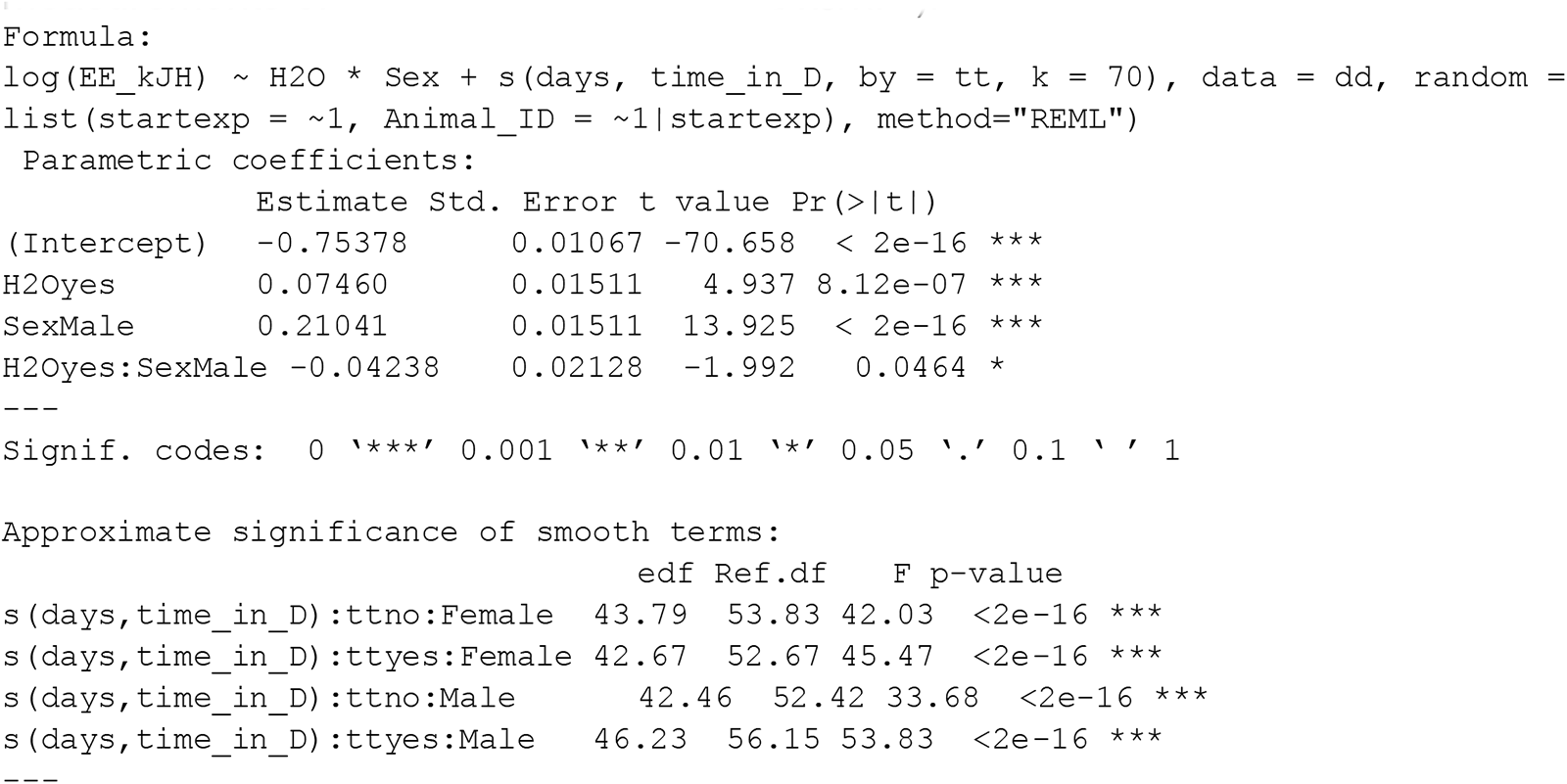
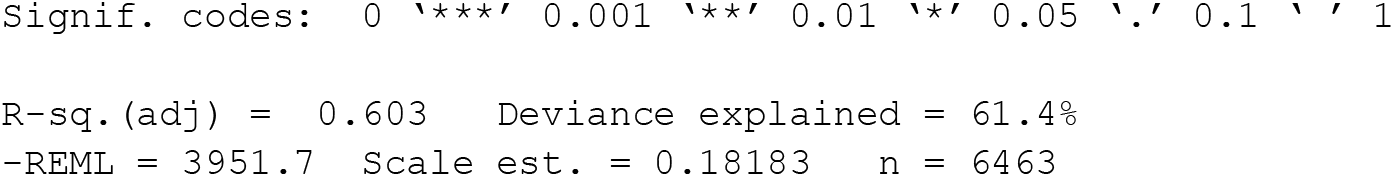
Generalized additive mixed models (GAMM) statical model and results for measurements of energy expenditure (EE kJ/hr_-1_).

**Table 3.**
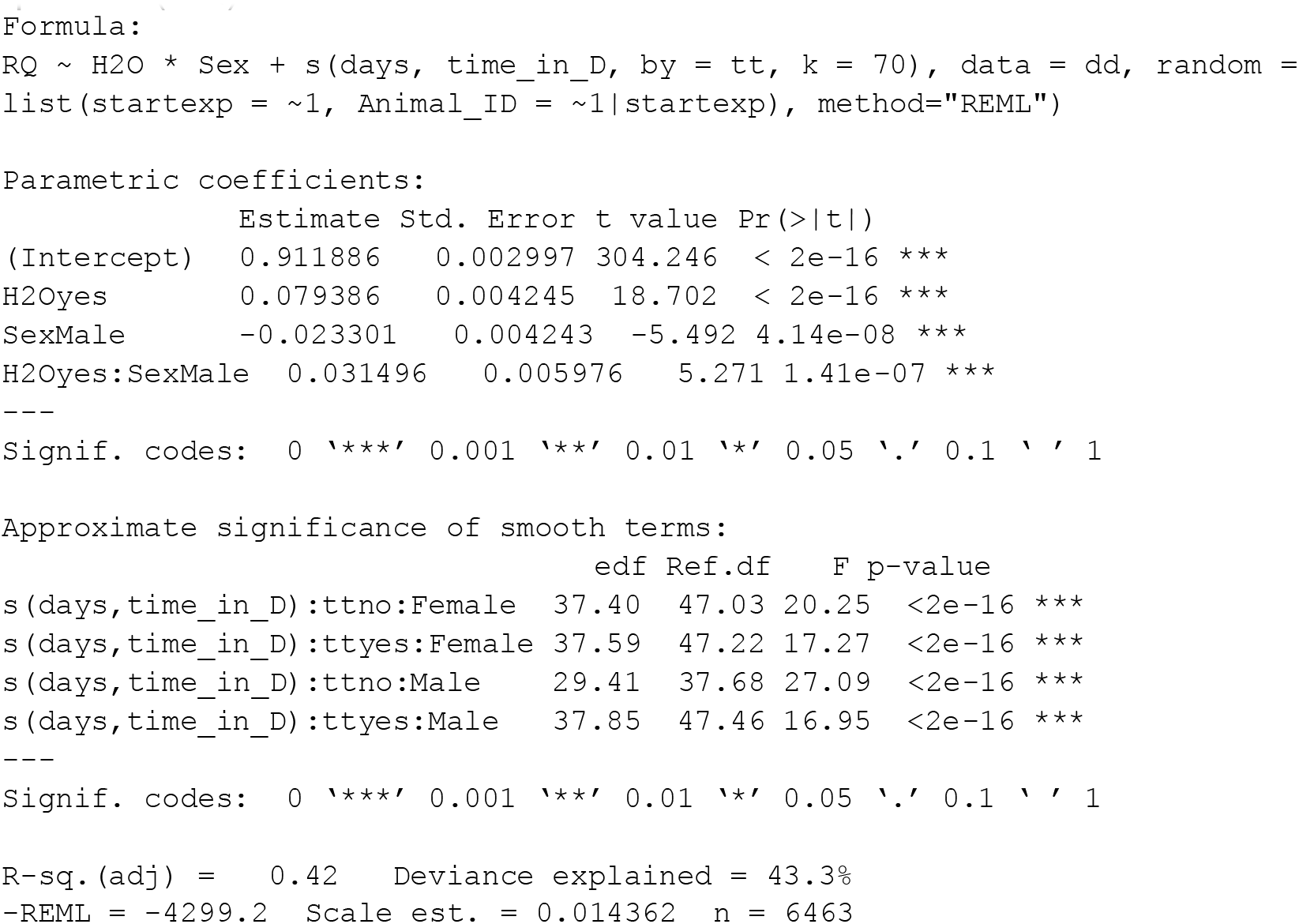
Generalized additive mixed models (GAMM) statical model and results for respiratory quotient (RQ).

